# AAV-mediated TBR1 replacement ameliorates behavioral and electrophysiological abnormalities in *Tbr1*-haploinsufficient mice

**DOI:** 10.64898/2026.09.22.753640

**Authors:** Tianshu Li, Zhuolei Jiao, Xiaqing Wang, Chengjie Yu, Yiting Yuan, Yuefang Zhang, Zilong Qiu

## Abstract

TBR1 haploinsufficiency causes a neurodevelopmental disorder associated with intellectual disability and autistic features, but the therapeutic potential of postnatal gene replacement remains incompletely defined. Here, we evaluated adeno-associated virus (AAV)-mediated human TBR1 replacement in *Tbr1*-haploinsufficient mice. A single retro-orbital injection of AAV-PHP.eB expressing human TBR1 under the human synapsin promoter was administered at postnatal day 3 at a dose of 2 × 10^11^ vector genomes. Compared with control-vector-treated mutants, treated mice exhibited increased TBR1 protein abundance and TBR1-, CTIP2-, and parvalbumin-positive cell counts in selected brain regions. Treatment improved sociability, social transmission of food preference, and buried-food seeking. Electroencephalographic (EEG) recordings showed attenuation of elevated relative theta power and improved theta suppression and gamma enhancement during social interaction. However, anterior commissure abnormalities persisted, correction of other EEG abnormalities was incomplete, and sleep–wake proportions showed no significant treatment improvement. These findings demonstrate that neonatal TBR1 replacement improves selected behavioral and electrophysiological outcomes despite persistent anatomical abnormalities. These findings support further development of gene replacement for TBR1 haploinsufficiency and highlight EEG measures as candidate biomarkers for monitoring treatment response.

## Introduction

Heterozygous pathogenic variants in TBR1 cause a neurodevelopmental disorder commonly characterized by intellectual disability, severe speech and language impairment, and autistic features. Affected individuals may also exhibit motor delay, hypotonia, and structural brain abnormalities, including reduced anterior commissure size and altered cortical or hippocampal organization. These findings implicate TBR1 dysfunction in both the anatomical development of the brain and the emergence of cognitive and behavioral impairments.^1^ The disease-associated variant spectrum includes protein-truncating variants, gene deletions, and missense substitutions. ^2^ Reduced functional TBR1 dosage provides a rationale for gene supplementation, although functional studies indicate that missense variants can differ in their effects on subcellular localization, transcriptional regulation, and interactions with other proteins.^2,3^

TBR1 is a T-box transcription factor expressed in postmitotic excitatory neurons and is essential for the development of forebrain circuits.^4^ During corticogenesis, TBR1 regulates the differentiation of early-born neurons, cortical organization, and the establishment of axonal projections.^4^ Its functions extend beyond initial neuronal specification: conditional deletion in cortical layer 6 neurons alters dendritic organization, synapse formation, and intrinsic electrophysiological properties, with abnormalities also occurring after loss of a single allele.^5^ In constitutive *Tbr1* heterozygous mice, impaired inter- and intra-amygdalar connectivity accompanies deficits in social interaction, communication, associative memory, and cognitive flexibility. ^6^ These animals also exhibit reduced neuronal activation in response to behavioral stimulation, linking altered developmental connectivity to abnormal circuit recruitment.^6^

The early developmental functions of TBR1 raise a central question for therapeutic intervention: can restoring its expression after birth improve brain function once circuit development has already been perturbed? Previous studies suggest that several consequences of *Tbr1* deficiency remain modifiable. Enhancing NMDA receptor activity through local amygdalar administration of D-cycloserine ameliorates behavioral deficits in *Tbr1* heterozygous mice.^6^ Pharmacological enhancement of WNT signaling also improves dendritic spine maturation and synaptic phenotypes in *Tbr1*-deficient models, and selected synaptic abnormalities remain responsive to lithium treatment in adulthood.^7,8^ These observations support the existence of a postnatal capacity for functional improvement and motivate direct evaluation of TBR1 supplementation.

Adeno-associated virus (AAV)-mediated gene replacement offers an approach to address the underlying dosage deficit and potentially influence multiple downstream processes regulated by TBR1. However, the outcome of replacement is likely to depend on the timing and cellular context of transgene expression. Ectopic *Tbr1* expression in embryonic cortical progenitors can disrupt neuronal migration, specification, and dendritic development, demonstrating that the consequences of TBR1 expression are context dependent.^9^ Evaluating a neuron-directed replacement strategy therefore requires assessment of functional outcomes alongside transgene expression and brain anatomy.

Measurements of neuronal population activity may help connect molecular intervention with behavioral improvement. Optogenetic stimulation of the basolateral amygdala in *Tbr1* heterozygous mice alters brain-wide activity patterns inferred from c-*fos* expression and modifies specific forms of social interaction, further implicating abnormal circuit recruitment in the behavioral phenotype.^10^ Electroencephalographic recordings aligned to social encounters provide a complementary approach for examining the temporal dynamics of neuronal activity during behavioral engagement. Combining these recordings with baseline EEG and behavioral assessments can help determine whether TBR1 replacement improves both spontaneous activity and the activity changes accompanying social interaction.

Here, we investigated the effects of neonatal TBR1 replacement in *Tbr1*-haploinsufficient mice using systemic delivery of AAV-PHP.eB encoding human TBR1 under the human synapsin promoter. Following administration at postnatal day 3, we evaluated behavioral outcomes, neuronal marker expression, anterior commissure anatomy, and baseline and social interaction– associated EEG activity. TBR1 replacement improved selected social behaviors and shifted social interaction–associated theta and gamma responses toward wild-type patterns in *Tbr1*-haploinsufficient mice, while anterior commissure abnormalities persisted. These findings provide preclinical evidence that early postnatal TBR1 replacement improves selected behavioral outcomes and highlight EEG measures as candidate biomarkers for monitoring treatment response.

## Results

### *Tbr1*-haploinsufficient mice exhibit reduced social investigation and increased locomotor activity

We first characterized heterozygous *Tbr1*-deficient (*Tbr1*^+/−^) mice carrying a deletion of exons 2–4 (Figure 1A). Immunoblotting demonstrated significantly reduced TBR1 protein abundance in heterozygous mice compared with wild-type (WT) controls, confirming the effect of the targeted deletion on TBR1 expression (Figures 1B and 1C).

**Figure 1.**
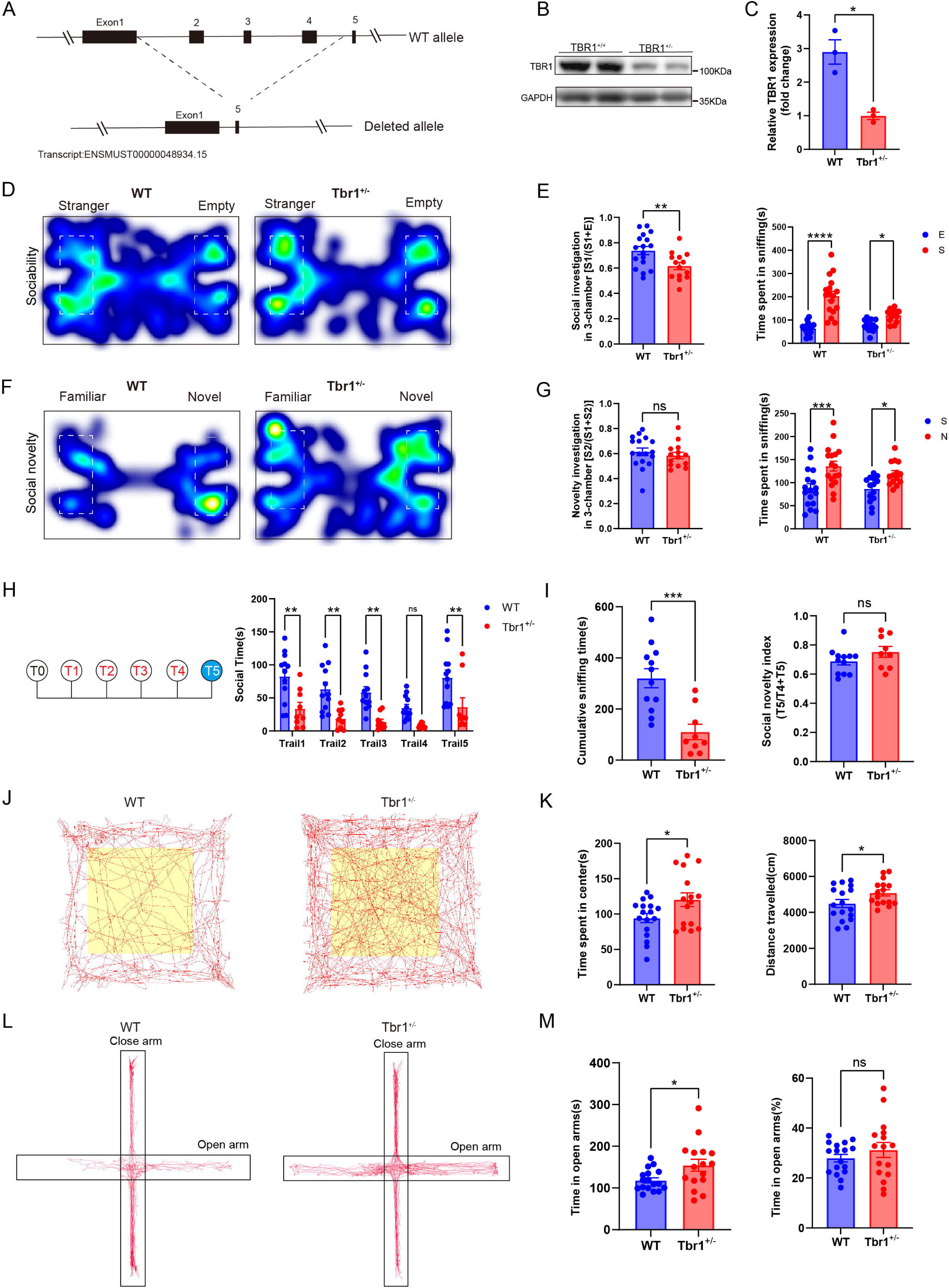
Tbr1-haploinsufficient mice exhibit reduced social investigation and increased locomotor activity. (A) Schematic of the wild-type (WT) and mutant *Tbr1* alleles, showing deletion of exons 2–4. Transcript: ENSMUST00000048934.15. (B and C) Representative immunoblots of TBR1 and GAPDH in the prefrontal cortex (PFC) from WT and *Tbr1*^+/−^ mice (B) and quantification of TBR1 protein abundance (C). GAPDH served as the loading control. TBR1 band intensities were normalized to GAPDH, and the resulting ratios were expressed as fold change relative to the mean value of the *Tbr1*^+/−^ group (set to 1). Each lane and each data point represent an independent biological sample. Group comparisons were performed using unpaired two-tailed t-tests with Welch’s correction (not assuming equal variances). (D and E) Representative heatmaps during the sociability phase of the three-chamber test (D), social investigation index (E, left), and time spent sniffing the stranger mouse or empty enclosure (E, right). The social investigation index was calculated as S1/(S1 + E), where S1 and E denote the time spent investigating the stranger mouse and empty enclosure, respectively. WT, n = 17; *Tbr1*^+/−^, n = 14. The social investigation index (E, left) was analyzed using an unpaired two-tailed t-test with Welch’s correction, and sniffing time (E, right) was analyzed using ordinary two-way ANOVA followed by Šídák’s multiple comparisons test. (F and G) Representative heatmaps during the social novelty phase (F), social novelty index (G, left), and time spent sniffing the familiar (F) and novel (N) mice (G, right). WT, n = 17; *Tbr1*^+/−^, n = 14. The social novelty index (G, left) was analyzed using an unpaired two-tailed t-test with Welch’s correction, and sniffing time (G, right) was analyzed using ordinary two-way ANOVA followed by Šídák’s multiple comparisons test. (H and I) Schematic of the repeated intruder test and social investigation time during each trial (H). The same intruder was presented during trials 1–4, followed by a novel intruder in trial 5. Cumulative sniffing time across the five trials and the social novelty index are shown in (I). The novelty index was calculated as T5/(T4 + T5), where T4 and T5 denote sniffing times during trials 4 and 5, respectively. WT, n = 12; *Tbr1*^+/−^, n = 9. Data in (H) were analyzed using ordinary two-way ANOVA followed by Šídák’s multiple comparisons test. Both cumulative sniffing time and the novelty index in (I) were analyzed using unpaired two-tailed t-tests with Welch’s correction. (J and K) Representative movement trajectories in the open-field test (J) and quantification of time spent in the center and total distance traveled (K). Yellow shading indicates the center zone. WT, n = 17; *Tbr1*^+/−^, n = 16. Data were analyzed using unpaired two-tailed t-tests with Welch’s correction. (L and M) Representative movement trajectories in the elevated plus maze (L) and quantification of absolute time and the proportion of time spent in the open arms (M). WT, n = 17; *Tbr1*^+/−^, n = 16. Data were analyzed using unpaired two-tailed t-tests with Welch’s correction. Data are presented as mean ±SEM. Each point represents an individual mouse. Mice were 6–8 weeks old, and both male and female mice were included. *p < 0.05, **p < 0.01, ***p < 0.001, and ****p < 0.0001; ns, not significant.

We next examined social behavior using the three-chamber test. During the sociability phase, both WT and *Tbr1*^+/−^ mice spent significantly more time investigating the unfamiliar mouse than the empty enclosure. However, the social investigation index was significantly lower in *Tbr1*^+/−^ mice, indicating attenuated sociability despite retained preference for a social stimulus (Figures 1D and 1E). During the subsequent social novelty phase, both genotypes spent more time investigating the novel mouse than the familiar mouse, and no significant genotype difference was detected in the social novelty index (Figures 1F and 1G).

To further assess social investigation across repeated encounters, we performed a repeated intruder test in which a new intruder was introduced during the fifth trial. Compared with WT controls, *Tbr1*^+/−^ mice exhibited significantly shorter social investigation times during trials 1–3 and trial 5, whereas the genotype difference during trial 4 was not significant (Figure 1H). Cumulative sniffing time across the five trials was also significantly reduced in *Tbr1*^+/−^ mice (Figure 1I). Nevertheless, the social novelty index calculated from trials 4 and 5 did not differ significantly between genotypes (Figure 1I). *Tbr1*^+/−^ mice exhibited reduced overall social investigation during free-contact encounters, without a significant genotype difference in the social novelty index.

We also assessed locomotor activity and exploration in the open-field and elevated plus-maze tests. In the open field, *Tbr1*^+/−^ mice traveled a greater total distance and spent more time in the center than WT controls (Figures 1J and 1K). In the elevated plus maze, heterozygous mice showed increased absolute time in the open arms, whereas the proportion of time spent in the open arms did not differ significantly between genotypes (Figures 1L and 1M). Thus, *Tbr1* deficiency was associated with increased locomotor activity and altered exploration. Together with the social assays, these results indicate that reduced social investigation in *Tbr1*^+/−^ mice occurs in the context of increased, rather than diminished, general activity.

### Neonatal AAV-mediated TBR1 replacement ameliorates neuronal marker abnormalities in *Tbr1*-haploinsufficient mice

To investigate whether neonatal TBR1 replacement could ameliorate cellular and anatomical abnormalities associated with *Tbr1* haploinsufficiency, we generated an AAV-PHP.eB vector expressing HA-tagged human TBR1 under the human synapsin promoter (Figure 2A). *Tbr1*^+/−^ mice received a retro-orbital injection of AAV-hTBR1 at postnatal day 3 (P3), at a total dose of 2 × 10^11^ vector genomes per mouse. WT and *Tbr1*^+/−^ mice receiving AAV-EGFP served as controls (Figure 2B, Figure S1A). Behavioral assessments began at P56, electrode implantation was performed at P120 for subsequent EEG recordings, and brain tissue was collected at P150 (Figure 2C).

**Figure 2.**
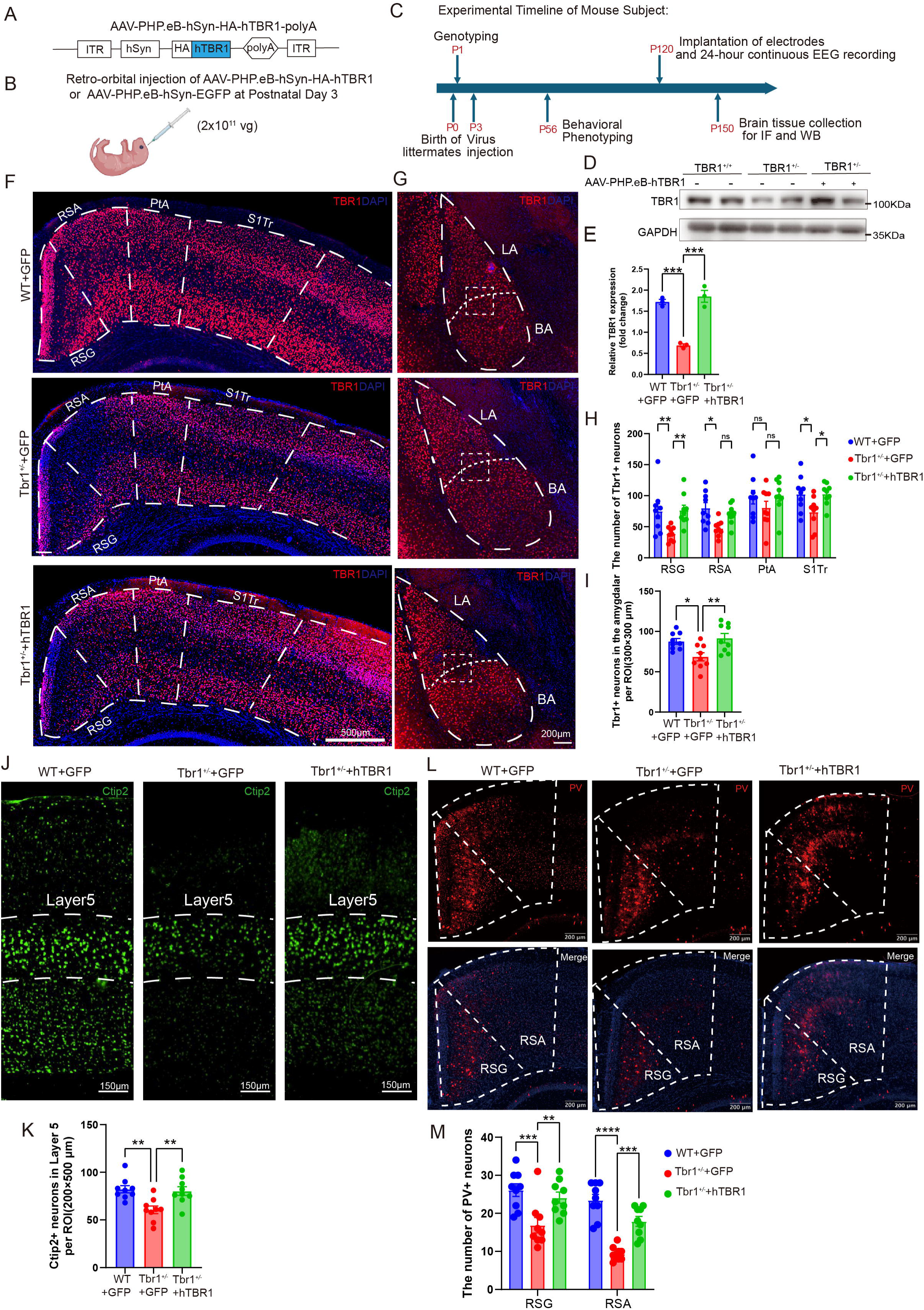
Neonatal AAV-mediated TBR1 replacement ameliorates neuronal marker abnormalities in *Tbr1*-haploinsufficient mice. (A) Schematic of the AAV-PHP.eB-hSyn-HA-hTBR1-polyA vector expressing HA-tagged human TBR1 under the human synapsin promoter. ITR, inverted terminal repeat; HA, hemagglutinin epitope tag; polyA, polyadenylation signal. (B) Schematic of retro-orbital administration of AAV-PHP.eB-hSyn-HA-hTBR1 or control AAV-PHP.eB-hSyn-EGFP at postnatal day 3 (P3), at a dose of 2 × 10¹¹ vector genomes per mouse. Experimental groups comprised wild-type mice receiving AAV-EGFP (WT+GFP), *Tbr1*^+/−^ mice receiving AAV-EGFP (*Tbr1*^+/−^+GFP), and *Tbr1*^+/−^ mice receiving AAV-hTBR1 (*Tbr1*^+/−^+hTBR1). (C) Experimental timeline showing genotyping at P1, vector administration at P3, behavioral phenotyping beginning at P56, electrode implantation at P120 for subsequent 24-h electroencephalographic recording, and brain tissue collection at P150 for immunofluorescence and immunoblot analyses. (D and E) Representative immunoblots (D) and quantification (E) of TBR1 protein expression in the prefrontal cortex (PFC). GAPDH served as a loading control. TBR1 band intensities were normalized to GAPDH, and the resulting ratios were expressed as fold change relative to the mean value of the *Tbr1*^+/−^+GFP group (set to 1). Each lane and each data point represent an independent biological sample; n = 3 mice per group. Group comparisons were performed using ordinary one-way ANOVA followed by Dunnett’s multiple comparisons test. (F) Representative immunofluorescence images showing TBR1 (red) and DAPI (blue) in the granular retrosplenial cortex (RSG), agranular retrosplenial cortex (RSA), parietal association cortex (PtA), and trunk region of the primary somatosensory cortex (S1Tr). Dashed outlines delineate the indicated cortical regions. (G) Representative immunofluorescence images showing TBR1 (red) and DAPI (blue) in the amygdala. LA, lateral amygdala; BA, basal amygdala. (H) Quantification of TBR1-positive cells in the RSG, RSA, PtA, and S1Tr. Each point represents one section (three sections per mouse; n = 3 mice per group). Data were analyzed using ordinary two-way ANOVA followed by Tukey’s multiple comparisons test. (I) Quantification of TBR1-positive cells in the amygdala per 300 × 300 μm regions of interest (ROI). Each point represents one section (three sections per mouse; n = 3 mice per group). Data were analyzed using ordinary one-way ANOVA followed by Dunnett’s multiple comparisons test. (J and K) Representative CTIP2 immunofluorescence images (green; J) and quantification of CTIP2-positive cells in cortical layer V per 200 × 500 μm ROI (K). Dashed lines in J delineate layer V in the S1Tr. Each point represents one section (three sections per mouse; n = 3 mice per group). Data were analyzed using ordinary one-way ANOVA followed by Dunnett’s multiple comparisons test. (L and M) Representative parvalbumin (PV) immunofluorescence images (L) and quantification of PV-positive cells in the RSG and RSA (M). Upper images show PV staining (red); lower images show merged PV and DAPI staining (blue). Dashed outlines delineate the indicated cortical regions. Each point represents one section (three sections per mouse; n = 3 mice per group). Data were analyzed using ordinary one-way ANOVA followed by Dunnett’s multiple comparisons test. Scale bars, 500 μm (F), 200 μm (G and L), and 150 μm (J). Data are presented as mean ±SEM. Both male and female mice were included Asterisks indicate significance for the comparisons marked by brackets: *p < 0.05, **p < 0.01, ***p < 0.001, and ****p < 0.0001.

Immunoblotting revealed significantly reduced TBR1 protein abundance in GFP-treated *Tbr1*^+/−^ mice compared with WT controls. AAV-hTBR1 administration significantly increased TBR1 abundance in mutant mice, with the group mean approaching that of WT controls (Figures 2D and 2E).

Regional immunostaining further demonstrated reduced TBR1-positive cell counts in the granular and agranular retrosplenial cortex (RSG and RSA), the trunk region of the primary somatosensory cortex (S1Tr), and the amygdala of GFP-treated mutants. TBR1 replacement significantly increased these counts in RSG, S1Tr, and the amygdala (Figures 2F–2I). Layer-specific analyses also revealed reduced TBR1-positive cell counts in cortical layers V and VI in GFP-treated mutants, both of which were significantly increased following treatment (Figures S1B-S1D).

We next examined whether TBR1 replacement affected additional neuronal markers. GFP-treated *Tbr1*^+/−^ mice exhibited fewer CTIP2-positive cells within cortical layer V than WT controls, and AAV-hTBR1 treatment significantly increased CTIP2-positive cell counts within the sampled regions (Figures 2J and 2K). Similarly, parvalbumin (PV)-positive cell counts were significantly reduced in both RSG and RSA in GFP-treated mutants and increased following TBR1 replacement (Figures 2L and 2M). These findings demonstrate that neonatal TBR1 replacement improves several neuronal developmental marker readouts beyond TBR1 immunoreactivity itself.

Despite these improvements in neuronal marker profiles, the anterior commissure abnormality persisted after treatment. The measured extent of the anterior commissure was markedly reduced in GFP-treated *Tbr1*^+/−^ mice compared with WT controls. This measurement did not differ significantly between GFP- and hTBR1-treated mutants, indicating no detectable improvement in the commissural defect under the treatment conditions examined (Figures S1E and S1F).

To assess changes in microglial abundance, we quantified IBA1-positive cells in RSG, RSA, and the parietal association cortex (PtA). No significant differences were detected between GFP-treated WT and mutant mice or between GFP- and hTBR1-treated mutants in these regions (Figures S1G and S1H).

### Neonatal TBR1 replacement selectively ameliorates behavioral abnormalities in *Tbr1*-haploinsufficient mice

We next examined whether neonatal TBR1 replacement improved behavioral outcomes in *Tbr1*^+/−^ mice. In the three-chamber sociability test, GFP-treated *Tbr1*^+/−^ mice exhibited a significantly lower social investigation index than WT controls, and this index was significantly increased following AAV-hTBR1 treatment (Figure 3A). No significant differences were detected in the social novelty index between GFP-treated WT and *Tbr1*^+/−^ mice or between the two *Tbr1*^+/−^ treatment groups (Figure 3B). During repeated intruder encounters, GFP-treated *Tbr1*^+/−^ mice showed reduced social investigation across all five trials compared with WT controls. TBR1 replacement significantly increased investigation in *Tbr1*^+/−^ mice during the first encounter, whereas treatment comparisons during trials 2–5 were not significant (Figure 3C). Cumulative sniffing time was numerically greater in AAV-hTBR1-treated *Tbr1*^+/−^ mice than in GFP-treated *Tbr1*^+/−^ mice (Figure 3D). Thus, treatment improved three-chamber sociability and initial social investigation, with more limited effects across repeated encounters.

**Figure 3.**
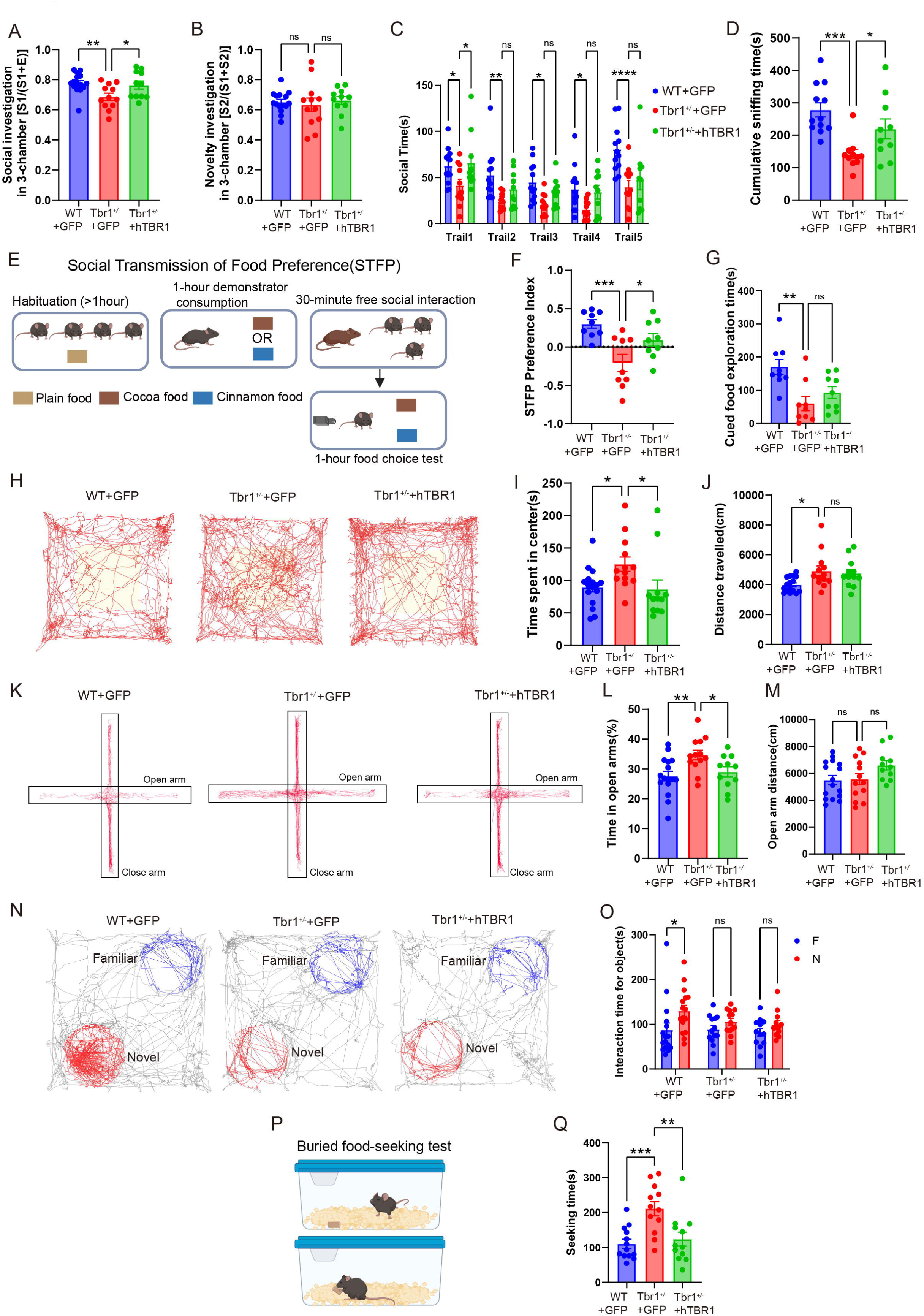
Neonatal AAV-mediated TBR1 replacement selectively ameliorates behavioral abnormalities in *Tbr1*-haploinsufficient mice. Wild-type mice received control AAV-EGFP (WT+GFP), and *Tbr1*^+/−^ mice received either AAV-EGFP (*Tbr1*^+/−^+GFP) or AAV-hTBR1 (*Tbr1*^+/−^+hTBR1). Vectors were administered by retro-orbital injection at postnatal day 3 at a dose of 2 × 10¹¹ vector genomes per mouse. (A) Social investigation index in the three-chamber sociability test, calculated as S1/(S1 + E), where S1 and E represent the time spent investigating the stranger mouse and empty enclosure, respectively. WT+GFP, n=15; *Tbr1*^+/−^+GFP, n=12; *Tbr1*^+/−^+hTBR1, n=11. Data were analyzed using ordinary one-way ANOVA followed by Dunnett’s multiple comparisons test. (B) Social novelty investigation index in the three-chamber social novelty test. WT+GFP, n=15; *Tbr1*^+/−^+GFP, n=12; *Tbr1*^+/−^+hTBR1, n=11. Data were analyzed using ordinary one-way ANOVA followed by Dunnett’s multiple comparisons test. (C) Social investigation time across five trials of the repeated social exposure test. The same stimulus mouse was presented in trials 1–4, followed by a novel stimulus mouse in trial 5. WT+GFP, n=12; *Tbr1*^+/−^+GFP, n=11; *Tbr1*^+/−^+hTBR1, n=10. Data were analyzed using ordinary two-way ANOVA followed by Dunnett’s multiple comparisons test. (D) Cumulative sniffing time across the five trials shown in C. WT+GFP, n=12; *Tbr1*^+/−^+GFP, n=11; *Tbr1*^+/−^+hTBR1, n=10. Data were analyzed using ordinary one-way ANOVA followed by Dunnett’s multiple comparisons test. (E) Schematic of the social transmission of food preference (STFP) test. Following habituation for more than 1 h, a demonstrator mouse consumed cocoa-or cinnamon-flavored food for 1 h and subsequently interacted with observer mice for 30 min. Observer mice underwent a 1-h choice test with both flavored foods. (F) STFP preference index WT+GFP, n=9; *Tbr1*^+/−^+GFP, n=9; *Tbr1*^+/−^+hTBR1, n=9. Data were analyzed using ordinary one-way ANOVA followed by Dunnett’s multiple comparisons test. The preference index was calculated as (cued food intake − non-cued food intake) / (cued food intake + non-cued food intake). (G) Time spent exploring the cued food Cued food denotes the flavor previously consumed by the demonstrator mouse. WT+GFP, n=9; *Tbr1*^+/−^+GFP, n=9; *Tbr1*^+/−^+hTBR1, n=9. Data were analyzed using ordinary one-way ANOVA followed by Dunnett’s multiple comparisons test (H, I and J) Representative movement trajectories (H) and quantification of time spent in the center (I) and total distance traveled (J) in the open-field test. The shaded area in H indicates the center zone. WT+GFP, n=16; *Tbr1*^+/−^+GFP, n=13; *Tbr1*^+/−^+hTBR1, n=12. Data were analyzed using ordinary one-way ANOVA followed by Dunnett’s multiple comparisons test. (K, L and M) Representative movement trajectories (K) and quantification of the proportion of time spent in the open arms (L) and distance traveled in the open arms (M) in the elevated plus-maze test. WT+GFP, n=16; *Tbr1*^+/−^+GFP, n=13; *Tbr1*^+/−^+hTBR1, n=11. Data were analyzed using ordinary one-way ANOVA followed by Dunnett’s multiple comparisons test. (N and O) Representative exploration trajectories (N) and interaction times with familiar and novel objects (O) in the novel object recognition test. Blue and red traces in K indicate exploration around the familiar and novel objects, respectively. F, familiar object; N, novel object. WT+GFP, n=16; *Tbr1*^+/−^+GFP, n=13; *Tbr1*^+/−^+hTBR1, n=12. Data were analyzed using ordinary two-way ANOVA followed by Šídák’s multiple comparisons test. (P and Q) Schematic of the buried food-seeking test (P) and quantification of the latency to locate the buried food (Q). WT+GFP, n=13; *Tbr1*^+/−^+GFP, n=12; *Tbr1*^+/−^+hTBR1, n=12. Data were analyzed using ordinary one-way ANOVA followed by Dunnett’s multiple comparisons test. Data are presented as mean ±SEM. Each point represents Each point represents an individual mouse. Both male and female mice were included. Asterisks indicate significance for the comparisons marked by brackets: *p < 0.05, **p < 0.01, ***p < 0.001, and ****p < 0.0001; ns, not significant.

We further assessed behavioral responses to socially transmitted food cues using the social transmission of food preference (STFP) assay (Figure 3E). GFP-treated *Tbr1*^+/−^ mice exhibited a reduced STFP preference index, shorter cued-food exploration time, and a lower cued-food exploration ratio than WT controls. TBR1 replacement significantly increased the preference index in *Tbr1*^+/−^ mice, whereas the increase in absolute cued-food exploration time was not statistically significant (Figures 3F and 3G).

Treatment also modified exploratory behavior. In the open-field test, GFP-treated mutants spent more time in the center and traveled a greater total distance than WT controls. TBR1 replacement significantly reduced center time in *Tbr1*^+/−^ mice, whereas total distance traveled did not differ significantly between the two mutant treatment groups (Figures 3H–3J). In the elevated plus maze, GFP-treated mutants spent a greater proportion of time in the open arms than WT controls, and this proportion was significantly reduced by TBR1 supplementation. Open-arm distance traveled did not differ significantly in either comparison (Figures 3K–3M). These findings indicate changes in exploratory behavior following treatment without a detectable reduction in open-field locomotor activity.

In the novel-object recognition assay, WT controls spent significantly more time investigating the novel object than the familiar object. A significant novel-object preference was not detected in either GFP- or hTBR1-treated *Tbr1*^+/−^ mice (Figures 3N and 3O). Therefore, the analysis did not demonstrate the emergence of a significant novel-object preference following treatment. In the buried food-seeking test, GFP-treated mutants required significantly longer than WT controls to locate the food. This latency was significantly reduced following TBR1 supplementation, demonstrating improved food-seeking performance (Figures 3P and 3Q).

Additional behavioral assessments revealed no significant differences in the number of marbles remaining unburied between GFP-treated WT and *Tbr1*^+/−^ mice or between the two mutant treatment groups (Figures S2A and S2B). Barnes-maze acquisition curves are presented in Figure S2C; in the subsequent preference assessment, all three groups spent significantly more time in the target region than in the opposite region (Figure S2D). Rotarod performance was broadly similar across groups, with no significant differences in the reported pairwise comparisons of latency to fall (Figures S2E–S2G).

### Neonatal TBR1 replacement reduces relative theta power and theta–beta coupling in *Tbr1*-haploinsufficient mice

To determine whether neonatal TBR1 replacement affected brain network activity, we performed continuous EEG recordings following electrode implantation, a 7-day recovery period, and 2 days of habituation (Figure 4A). Analysis of baseline EEG revealed significantly increased relative theta-band power (4–8 Hz) in GFP-treated *Tbr1*^+/−^ mice compared with WT controls. AAV-hTBR1 treatment significantly reduced relative theta power in *Tbr1*^+/−^ mice (Figures 4B and 4C). GFP-treated mutants also exhibited reduced relative delta power and increased relative gamma power, neither of which changed significantly following treatment (Figures 4D and 4E). Relative alpha and beta power did not differ significantly between GFP-treated WT and *Tbr1*^+/−^ mice or between the two mutant treatment groups (Figures 4F and 4G). The supplementary band-power analysis likewise identified lower delta and higher gamma power in GFP-treated *Tbr1*^+/−^ mice, with no significant treatment effects on any of the five absolute band-power measures (Figures S3A–S3E).

**Figure 4.**
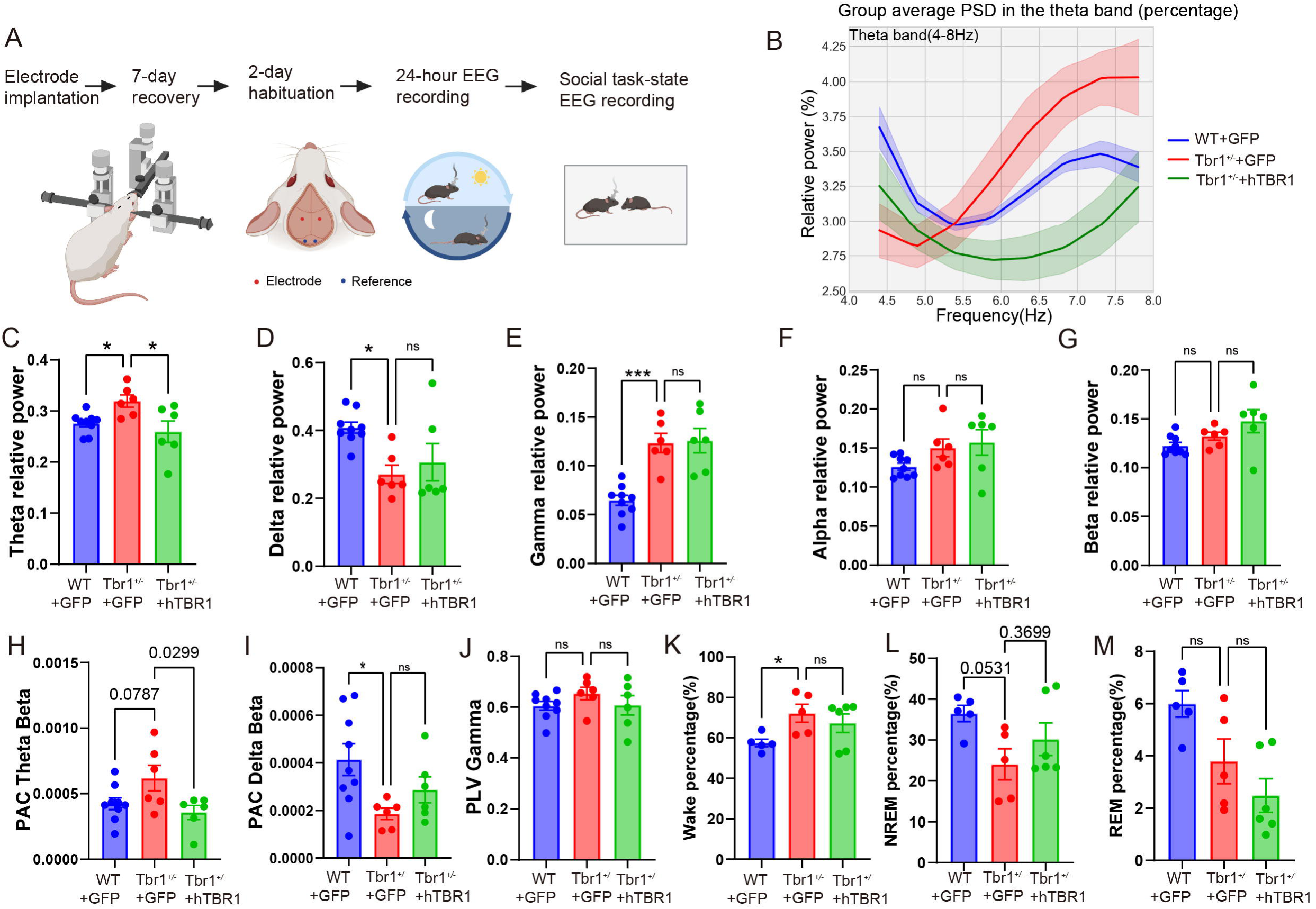
Neonatal AAV-mediated TBR1 replacement reduces relative theta power and theta–beta coupling in *Tbr1*-haploinsufficient mice. EEG recordings were obtained from wild-type mice receiving control AAV-EGFP (WT+GFP), *Tbr1*^+/−^ mice receiving AAV-EGFP (*Tbr1*^+/−^+GFP), and *Tbr1*^+/−^ mice receiving AAV-hTBR1 (*Tbr1*^+/−^+hTBR1) following neonatal vector administration. (A) Schematic of electrode placement and the electroencephalographic (EEG) recording protocol. Electrode implantation was followed by a 7-day recovery period, 2 days of habituation, and 24-h continuous EEG recording. Subsequent EEG recordings during social interaction were performed as illustrated. Red and blue dots indicate recording and reference electrodes, respectively. (B) Group-averaged relative power spectra within the theta band (4–8 Hz). Solid lines indicate group means, and shaded areas indicate SD. (C–G) Quantification of relative EEG power in the theta (C), delta (D), gamma (E), alpha (F), and beta (G) frequency bands. Baseline EEG analyses were performed on the first night segment of the 24-h continuous recording, using the first two ECoG channels, across all vigilance states within that segment. Relative power was calculated as the power in each band normalized to the total power in the 1–100 Hz range. Analyses. WT+GFP, n=9; *Tbr1*^+/−^+GFP, n=6; *Tbr1*^+/−^+hTBR1, n=6. Data were analyzed using ordinary one-way ANOVA followed by Dunnett’s multiple comparisons test. (H and I) Quantification of theta–beta (H) and delta–beta (I) phase–amplitude coupling (PAC). PAC was quantified using the modulation index (MI), with theta (4–8 Hz) and delta (1–4 Hz) as the phase-providing bands and beta (13–30 Hz) as the amplitude-providing band. WT+GFP, n=9; *Tbr1*^+/−^+GFP, n=6; *Tbr1*^+/−^+hTBR1, n=6.Data were analyzed using ordinary one-way ANOVA followed by Dunnett’s multiple comparisons test. (J) Gamma-band phase-locking value (PLV) between analyzed channel pairs. WT+GFP, n=9; *Tbr1*^+/−^+GFP, n=6; *Tbr1*^+/−^+hTBR1, n=6. Data were analyzed using ordinary one-way ANOVA followed by Dunnett’s multiple comparisons test. (K–M) Percentage of analyzed recording time classified as wakefulness (K), non-rapid eye movement (NREM) sleep (L), and rapid eye movement (REM) sleep (M). Sleep–wake states were classified using standard criteria based on EEG and electromyographic (EMG) activity, with a 2.5-s epoch duration. WT+GFP, n=5; *Tbr1*^+/−^+GFP, n=5; *Tbr1*^+/−^+hTBR1, n=6. Data were analyzed using ordinary one-way ANOVA followed by Dunnett’s multiple comparisons test. Blue, red, and green denote WT+GFP, *Tbr1*^+/−^+GFP, and *Tbr1*^+/−^+hTBR1 groups, respectively. Bar graphs show mean ±SEM. Each point represents an individual mouse. Both male and female mice were included. Asterisks indicate significance for the comparisons marked by brackets: *p < 0.05 and ***p < 0.001; ns, not significant.

We next examined cross-frequency interactions using phase–amplitude coupling (PAC). Theta– beta PAC was numerically higher in GFP-treated *Tbr1*^+/−^ mice than in WT controls, although this difference did not reach statistical significance (P = 0.0787). TBR1 replacement significantly reduced theta–beta PAC relative to GFP-treated *Tbr1*^+/−^ mice (P = 0.0299; Figure 4H). Delta– beta PAC was significantly lower in GFP-treated *Tbr1*^+/−^ mice than in WT controls and was not significantly altered by treatment (Figure 4I). Additional analyses revealed increased alpha–beta PAC in GFP-treated *Tbr1*^+/−^ mice without a significant treatment effect, whereas delta–gamma PAC did not differ significantly in either comparison (Figures S3F and S3G).

Measures of oscillatory synchronization showed no significant differences in gamma-band phase-locking values or theta- and delta-band coherence between GFP-treated WT and *Tbr1*^+/−^ mice or between the two mutant treatment groups (Figure 4J; Figures S3H and S3I). Gamma bandwidth also did not differ significantly in these comparisons (Figure S3J). However, beta-to-gamma and theta-to-gamma power ratios were significantly lower in GFP-treated mutants than in WT controls, and neither ratio was significantly altered by TBR1 replacement (Figures S3K and S3L).

Finally, we assessed the distribution of wakefulness and sleep states during EEG monitoring. GFP-treated *Tbr1*^+/−^ mice spent a significantly greater proportion of recording time awake than WT controls (Figure 4K). The proportion of non-rapid eye movement (NREM) sleep was numerically lower in *Tbr1*^+/−^ mice, but the difference was not statistically significant (P = 0.0531; Figure 4L). Rapid eye movement (REM) sleep proportions also did not differ significantly between GFP-treated WT and *Tbr1*^+/−^ mice (Figure 4M). TBR1 replacement did not significantly alter the proportions of wakefulness, NREM sleep, or REM sleep relative to GFP-treated *Tbr1*^+/−^ mice (Figures 4K–4M). Representative temporal distributions of the classified states are shown in Figure S3M.

### Neonatal TBR1 replacement improves theta and gamma modulation during social interaction

To investigate the relationship between EEG activity and social behavior, we first examined the association between theta power and social interaction time. Theta power was negatively correlated with social interaction time (Pearson’s r = −0.5563, P < 0.01; Figure 5A), linking higher theta power to reduced social engagement. We subsequently recorded EEG during encounters with unfamiliar mice and aligned the recordings to social-contact onset, examining activity within 2.5 s before and after contact (Figure 5B).

**Figure 5.**
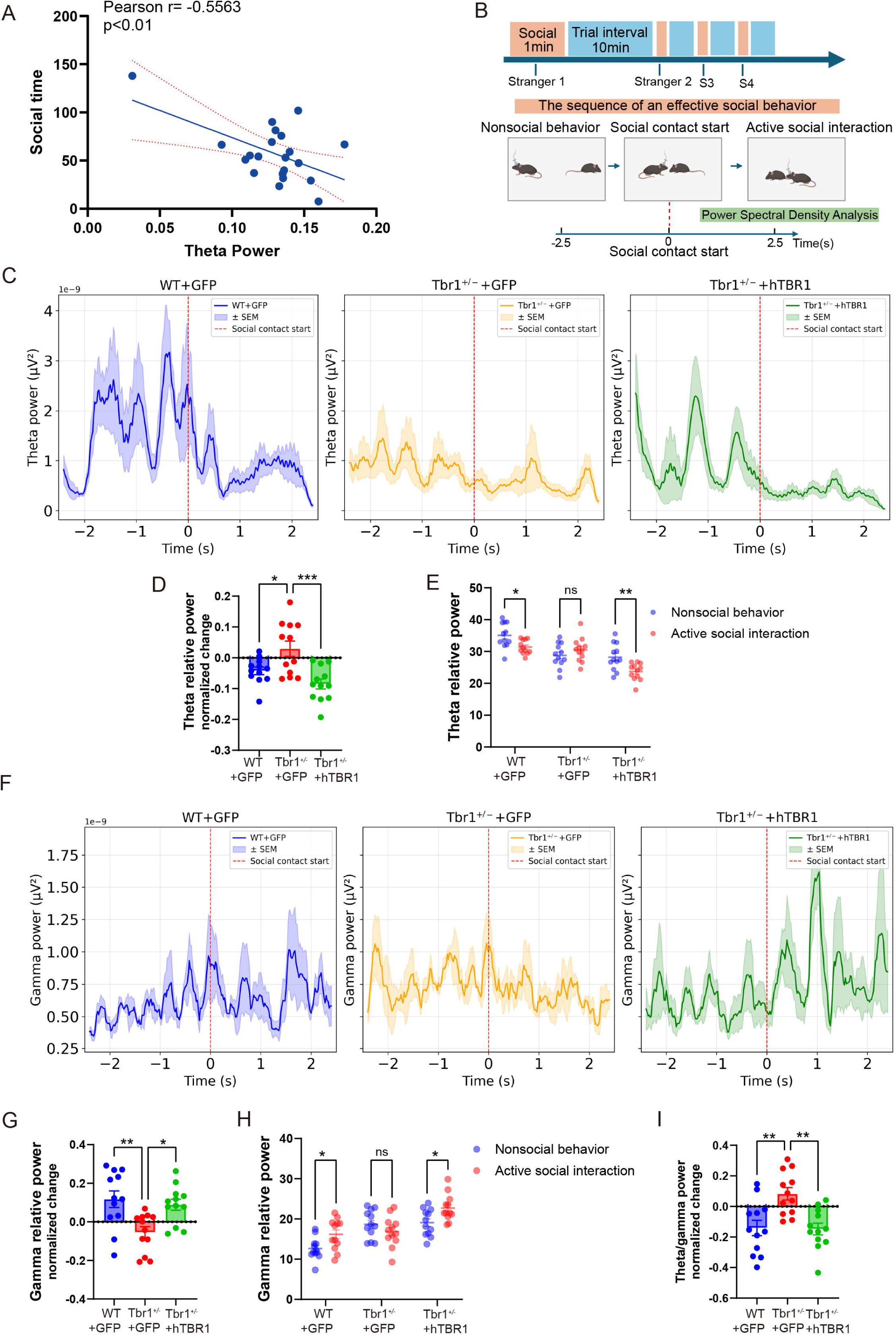
Neonatal AAV-mediated TBR1 replacement improves theta and gamma modulation during social interaction in *Tbr1*-haploinsufficient mice. (A) Correlation between theta power and social interaction time (Pearson’s r = −0.5563, p < 0.01). The solid line indicates the linear regression fit, and dotted lines indicate 95% confidence intervals of the linear regression. Theta power was derived from the 24-h continuous EEG recordings analyzed in Supplementary Figure 3(B), and social interaction time was obtained from trial 1 of the repeated intruder test in Figure 3(C). Data from all three experimental groups (WT+GFP, n=9; *Tbr1*^+/−^+GFP, n=6; *Tbr1*^+/−^+hTBR1, n=6) were pooled for this analysis. Each point represents an individual mouse. Data were analyzed using Pearson correlation. (B) Schematic of EEG recording during social interaction. Successive unfamiliar mice were presented in 1-min trials separated by 10-min intervals. EEG segments were aligned to social-contact onset (time 0), encompassing 2.5 s before and 2.5 s after contact. The schematic illustrates the transition from nonsocial behavior to active social interaction. (C) Mean theta-power time courses aligned to social-contact onset in wild-type mice receiving control AAV-EGFP (WT+GFP), *Tbr1*^+/−^ mice receiving AAV-EGFP (*Tbr1*^+/−^+GFP), and *Tbr1*^+/−^ mice receiving AAV-hTBR1 (*Tbr1*^+/−^+hTBR1). Shaded areas indicate ±SD, and red dashed lines mark social-contact onset. (D) Normalized change in relative theta power associated with the transition from nonsocial behavior to active social interaction. Each point represents a social-interaction event, with 12 events analyzed per group (WT+GFP, n = 3; *Tbr1*^+/−^+GFP, n=2 and *Tbr1*^+/−^+hTBR1 mice, n=3). Data were analyzed using ordinary one-way ANOVA followed by Dunnett’s multiple comparisons test. (E) Relative theta power during nonsocial behavior and active social interaction within each experimental group. Blue and pink points indicate nonsocial and social epochs, respectively. Each point represents a social-interaction event, with 12 events analyzed per group (WT+GFP, n = 3; *Tbr1*^+/−^+GFP, n = 2 and *Tbr1*^+/−^+hTBR1 mice, n = 3). Data were analyzed using ordinary two-way ANOVA followed by Šídák’s multiple comparisons test. (F) Mean gamma-power time courses aligned to social-contact onset in the three experimental groups. Shaded areas indicate ±SD, and red dashed lines mark social-contact onset. (G) Normalized change in relative gamma power associated with the transition from nonsocial behavior to active social interaction. Each point represents a social-interaction event, with 12 events analyzed per group (WT+GFP, n = 3; *Tbr1*^+/−^+GFP, n = 2 and *Tbr1*^+/−^+hTBR1 mice, n = 3). Data were analyzed using ordinary one-way ANOVA followed by Dunnett’s multiple comparisons test. (H) Relative gamma power during nonsocial behavior and active social interaction within each experimental group. Colors indicate behavioral conditions as in E. Each point represents a social- interaction event, with 12 events analyzed per group (WT+GFP, n = 3; *Tbr1*^+/−^+GFP, n = 2 and *Tbr1*^+/−^+hTBR1 mice, n = 3). Data were analyzed using ordinary two-way ANOVA followed by Šídák’s multiple comparisons test. (I) Normalized change in the theta-to-gamma power ratio associated with social interaction. Horizontal dashed lines in D, G, and I indicate no change. Each point represents a social-interaction event, with 12 events analyzed per group(WT+GFP, n = 3; *Tbr1*^+/−^+GFP, n = 2 and *Tbr1*^+/−^+hTBR1 mice, n = 3). Data were analyzed using ordinary one-way ANOVA followed by Dunnett’s multiple comparisons test. Relative power was calculated as the power in each frequency band divided by the total power in the 1–100 Hz range (expressed as a percentage). Normalized changes in D, G, and I were calculated as (social − nonsocial) / (social + nonsocial). Theta and gamma frequency bands were 4–8 Hz and 30–80 Hz, respectively. Bar graphs show mean ± SEM, and horizontal summary lines in E and H indicate the mean. For D, E, G, H, and I, all mice were female. Asterisks indicate significance for the comparisons marked by brackets: *p < 0.05, **p < 0.01, and ***p < 0.001; ns, not significant.

Social-contact-aligned recordings revealed distinct patterns of theta modulation across groups (Figure 5C; Figure S4A). In WT controls, relative theta power decreased significantly during active social interaction compared with preceding nonsocial behavior (Figures 5D and 5E). This decrease was not significant in GFP-treated *Tbr1*^+/−^ mice, whereas hTBR1-treated *Tbr1*^+/−^ mice exhibited a significant interaction-associated reduction in relative theta power (Figures 5D and 5E). Between-group comparisons of normalized changes in relative theta power further demonstrated impaired theta modulation in GFP-treated mutants and a significant shift toward theta suppression following TBR1 replacement (Figures 5D).

Gamma activity exhibited a complementary pattern around social-contact onset (Figure 5F; Figure S4B). Relative gamma power increased significantly during active social interaction in WT controls, whereas no significant increase was detected in GFP-treated *Tbr1*^+/−^ mice. A significant interaction-associated increase was observed in hTBR1-treated *Tbr1*^+/−^ mice (Figures 5G and 5H). Consistent with these within-group comparisons, the normalized change in relative gamma power was significantly lower in GFP-treated *Tbr1*^+/−^ mice than in WT controls and was significantly increased following TBR1 replacement (Figure 5G). The normalized change in the theta-to-gamma power ratio also differed significantly between WT controls and GFP-treated *Tbr1*^+/−^ mice. Treatment shifted this measure in the negative direction, consistent with the accompanying reduction in relative theta power and increase in relative gamma power during social interaction (Figure 5I).

Additional frequency-band analyses identified a significant treatment effect on delta modulation: hTBR1-treated *Tbr1*^+/−^ mice exhibited a greater normalized change in relative delta power than GFP-treated *Tbr1*^+/−^ mice. However, the corresponding comparison between GFP-treated WT and *Tbr1*^+/−^ mice was not significant (Figure S4C). Normalized changes in relative beta and alpha power did not differ significantly between GFP-treated WT and *Tbr1*^+/−^ mice or between the two mutant treatment groups (Figures S4D and S4E).

## Discussion

This study demonstrates that neonatal AAV-mediated replacement of human TBR1 improves selected cellular, behavioral, and electrophysiological outcomes in *Tbr1*-haploinsufficient mice. Treatment increased TBR1 abundance and several neuronal marker-positive cell counts, improved sociability and food-related behavioral performance, and modified EEG responses during social interaction. These benefits coexisted with a persistent anterior commissure abnormality, incomplete correction of baseline EEG abnormalities, and no significant improvement in sleep–wake measures. The findings support the therapeutic potential of augmenting TBR1 expression during early postnatal development while identifying limits to the responsiveness of different disease-associated outcomes.

The rationale for replacement extends beyond the role of TBR1 in initial brain patterning. TBR1 controls the differentiation of early-born cortical neurons and contributes to postnatal neuronal identity, dendritic development, and synaptic connectivity.^4,5^ Previous studies have demonstrated that selected consequences of *Tbr1* deficiency remain modifiable: NMDA receptor modulation improved behavioral outcomes in constitutive heterozygotes, and WNT-associated interventions restored synaptic phenotypes in several *Tbr1* mutant models, including after treatment in adulthood.^6,7,8^ Earlier work also reported rescue of axonal defects through wild-type *Tbr1* expression in cultured neurons.^6^ Our study extends this evidence by combining systemic neonatal delivery of human TBR1 with later assessments of behavior, regional marker expression, and baseline and social interaction-associated EEG activity. These complementary outcomes support continued investigation of gene replacement as an intervention for TBR1 haploinsufficiency.

The dissociation between functional improvement and persistent anterior commissure abnormalities is particularly informative. Commissural abnormalities have been reported in *Tbr1*-deficient mice and in individuals with TBR1-related neurodevelopmental disorder.^1,6^ In the present study, improvement in selected behaviors and EEG responses occurred without detectable correction of the measured commissural defect. This pattern is consistent with the possibility that neuronal signaling and circuit recruitment remain modifiable even when a major anatomical abnormality persists. It does not establish that the anterior commissure is dispensable for the affected behaviors. Failure to improve this structure could reflect an earlier developmental requirement for TBR1, delayed onset of transgene expression, or insufficient targeting of the relevant projection neurons. Comparisons across treatment ages and direct axonal tracing will be needed to distinguish these explanations.

The histological findings suggest that gene replacement influences neuronal phenotypes beyond increasing TBR1 immunoreactivity. Increased CTIP2-positive labeling in cortical layer V and PV-positive labeling in retrosplenial cortex could reflect improved neuronal differentiation, maintenance of marker expression, or indirect changes in circuit activity. However, marker-positive cell counts cannot distinguish these possibilities from changes in cell survival or abundance. Moreover, an increase in TBR1-positive cells after transgene delivery does not demonstrate restoration of endogenous cell identities, because expression may extend to neurons that normally express little TBR1. The accompanying PV changes provide a rationale for examining inhibitory circuit function, but neither these counts nor the EEG results establish correction of excitation–inhibition balance. Cell-type-resolved transcriptional profiling and direct measurements of excitatory and inhibitory synaptic function would help identify which downstream processes mediate benefit.

Behavioral improvement was selective. The clearest social effects were an increased three-chamber sociability index and greater investigation during the first intruder encounter; cumulative investigation across repeated encounters did not show a statistically significant treatment effect. Social novelty indices showed no established baseline deficit to rescue. Improvements in social transmission of food preference and buried-food seeking broaden the evidence for functional benefit, although these tasks depend on sensory processing, motivation, and investigation as well as learning. Changes in center and open-arm occupancy also require interpretation alongside locomotor activity, which was not significantly reduced by treatment. The behavioral profile therefore supports improvement in specific functions rather than generalized normalization of social behavior, cognition, or anxiety.

The EEG findings similarly indicate selective responsiveness. TBR1 replacement reduced the elevated relative theta power observed in GFP-treated mutants, whereas several other spectral abnormalities and the increased wake fraction did not show significant improvement. Treatment also reduced theta–beta phase–amplitude coupling, although the corresponding untreated genotype difference was not statistically significant. This coupling result therefore demonstrates modulation by treatment without establishing correction of a pre-existing abnormality. Relative power describes the distribution of activity across frequency bands and cannot alone establish a reduction in absolute theta activity. In addition, differences in sleep–wake composition may influence spectra aggregated across prolonged recordings. Analyses within matched vigilance states will be important for defining the physiological basis of the baseline EEG effects.

The social interaction recordings add a dynamic dimension to these observations. WT mice showed a reduction in relative theta power and an increase in relative gamma power during active social interaction. These responses were attenuated in GFP-treated mutants and improved following TBR1 supplementation, with significant treatment effects on the normalized modulation measures. The recovery of task-associated gamma modulation despite the absence of a significant treatment effect on elevated baseline relative gamma power suggests that tonic spectral composition and responsiveness to social events represent partly separable aspects of network function. Prior work showing that optogenetic theta-burst stimulation of the basolateral amygdala changes selected social behaviors in *Tbr1*-deficient mice provides independent evidence that activity within persisting circuits can influence behavioral output.^10^ The present EEG measurements, however, do not localize the circuits responsible for the treatment response.

The inverse association between theta power and social interaction time further supports investigation of EEG measures as candidate indicators of functional impairment and treatment response. This association remains exploratory: it does not establish that elevated theta activity causes reduced social engagement, and a relationship across pooled groups may differ from a relationship within each genotype or treatment group. Event-associated spectral changes may also reflect movement, sniffing, or arousal accompanying social contact. Validation will therefore require longitudinal measurements, analyses accounting for repeated encounters within individual animals, and comparisons that control these behavioral variables. Whether related EEG features occur in individuals with TBR1-related disorder also remains to be established before these mouse measures can be considered translational biomarkers.

Expression control is a central consideration for further development. The hSyn-driven cassette is designed to direct neuronal expression but does not reproduce the endogenous spatial distribution or regulation of TBR1. Consequently, a tissue-level protein measurement approaching the WT mean may conceal substantial variation among cell types or individual neurons. The importance of cellular context is supported by experiments in which embryonic *Tbr1* misexpression in cortical progenitors altered neuronal migration, differentiation, dendritic development, and callosal projections.^9^ Those experiments do not define a toxicity threshold for neonatal replacement, but they support evaluating expression level and cellular targeting explicitly. Dose-ranging studies, transgene mapping, and a WT group receiving the therapeutic vector would help define the therapeutic margin. No significant differences in IBA1-positive cell counts were detected in the sampled regions, but this limited observation does not establish comprehensive or long-term safety.

Delivery also requires separate translational validation. The enhanced CNS entry of AAV-PHP.eB depends on interactions with LY6A in permissive mouse strains.^11^ The related AAV-PHP.B capsid did not reproduce its enhanced mouse CNS tropism in the tested adult rhesus macaques, underscoring the limits of extrapolating delivery across species.^12^ Further development will require a capsid and administration route that achieve appropriate exposure in relevant human neuronal populations, together with assessment of peripheral distribution, immune responses, and dose-dependent toxicity.

The treatment window and genetic scope of the findings are equally important. Administration at P3 preceded behavioral assessment, so the present design cannot distinguish prevention or attenuation of emerging abnormalities from reversal of established symptoms. Juvenile and adult treatment cohorts are needed to define whether benefit remains achievable after clinically relevant deficits have developed. Efficacy was also evaluated in a haploinsufficiency model, whereas patient-associated TBR1 variants can produce differing effects on localization, transcriptional regulation, and protein interactions.^2,3^ Gene replacement has the clearest rationale when reduced functional dosage is the principal mechanism; its suitability for individual missense alleles will require variant-specific testing. Patient-derived neuronal models could help determine whether added WT TBR1 corrects the relevant molecular and physiological defects.

These results support further development of TBR1 gene replacement with behavioral and electrophysiological outcomes assessed alongside anatomical measures. Defining the effective treatment window, the required cellular distribution of expression, and a clinically relevant delivery strategy will determine whether the functional benefits observed after neonatal intervention can be extended to a broader therapeutic setting.

## Materials and methods

### Mice

All animal experiments were approved by the Institutional Animal Care and Use Committee of Songjiang Hospital Affiliated to Shanghai Jiao Tong University School of Medicine (ACE-004-2025). *Tbr1*^+/−^ and WT littermates on a C57BL/6J background were housed under SPF conditions (12-h light/dark cycle, lights on 7:00, off 19:00) with ad libitum food and water. Both sexes were included. The Tbr1 allele was generated by CRISPR/Cas9-mediated deletion (gB1A1–gB2A2, 1895 bp), confirmed by Sanger sequencing. Genotyping PCR used primers flanking the deleted region: forward, 5′-ATGTGGAAGGGATAATAACCACCG-3′; reverse, 5′- TTTAGCCCACCAGTTTCCGAGATT-3′. Products of 2296 bp and 401 bp corresponded to WT and deleted alleles, respectively.

### AAV vector construction and injection

AAV-PHP.eB vectors expressing hSyn-HA-hTBR1 or hSyn-GFP were packaged by PackGene Inc. All constructs were verified by sequencing. AAV-PHP.eB was injected via the retro-orbital route at P3 (20 μl, 1 × 10^13^ vg/ml). Mice were anesthetized on ice for approximately 3 min and placed on one side. Injection was performed perpendicular to the table at the 6 o’clock position of the eye at a depth of approximately 1 mm. The needle was slowly withdrawn after 3–5 sec. Mice were recovered on a 37 °C heating pad and returned to the cage with nesting material.

### Behavioral testing

Behavioral testing began at P56. Mice were handled for 3 consecutive days before testing and acclimated to the testing room for at least 30 min before each test. Both male and female mice were used, and sex was balanced across groups. Tests were performed from low to high stress in the following order: open field, elevated plus maze, novel object recognition, three-chamber social test, repeated social interaction, marble burying, social transmission of food preference (STFP), food burying, Barnes maze, and rotarod, with an inter-test interval of 24-48 h. Unless otherwise stated, behavior was recorded and analyzed using EthoVision XT (Noldus) and Python-based code, with scoring performed by observers blinded to genotype and treatment.

#### Three-chamber social test

The apparatus consisted of left, middle, and right chambers. Mice were habituated to the empty apparatus for 10 min on the day before testing and on the test day. In the social preference phase, a same-sex stranger mouse was placed under a wire cup in one side chamber, with an empty cup in the other; the test mouse was placed in the middle chamber, and interaction time was recorded for 10 min. In the social novelty phase, the empty cup was replaced with a second stranger mouse, and interaction was recorded for another 10 min.

#### Repeated social interaction test

Mice underwent four 3-min direct contact sessions with the same stranger mouse, separated by 5 min intervals, followed by a fifth 3-min session with a new stranger mouse. All stimulus mice were same-sex. Sniffing and close social interaction time were recorded and analyzed by a blinded observer using MATLAB.

#### Open field test

Mice were placed in the center of a 40 cm × 40 cm square arena under approximately 50 lux illumination and allowed to explore freely for 10 min. Total distance traveled, mean speed, speed per minute, and time spent in the central zone (20 cm × 20 cm) were recorded and analyzed.

#### Elevated plus maze

The open and closed arms were 67.2 cm (length) × 6 cm (width) × 63 cm (height). Mice were placed in the center of the maze with the head facing an open arm and allowed to explore freely for 10 min. Cumulative time spent in open and closed arms and movement trajectories were analyzed.

#### Novel object recognition

Mice were placed in a 40 cm × 40 cm square box. The test included habituation, familiarization with identical objects, and novel object discrimination, with 2-h intervals between stages. During familiarization, two identical cylinders (object 1) were placed in two opposite corners, and mice explored freely for 10 min; during discrimination, one cylinder (object 1) and one cone (object 2) were placed in the two corners, and mice explored freely for 10 min.

#### Social transmission of food preference (STFP)

On day 1, pellet food was removed from all cages to initiate food deprivation. On day 2, two food jars containing powdered chow were placed in each cage, and food intake was monitored hourly until each mouse consumed approximately 0.2 g. One WT “demonstrator” mouse per cage was then singly housed and food-deprived for 18 h with water available. On day 3, demonstrators received flavored chow (cocoa or cinnamon) for 1 h; meanwhile, food and water were removed from observer cages. Demonstrators were placed into observer cages for 30 min, and nose contacts were recorded. Observer mice then underwent 18 h of food deprivation, singly housed with water only. On day 4, each observer was presented with the cue flavor and a novel flavor for 1 h, and intake, jar approaches, and feeding behavior were recorded. Preference index = (cue food intake − novel food intake) / (cue food intake + novel food intake).

#### Buried food seeking test

Food was removed from the home cage 24 h before testing, residual crumbs were cleared, and water was retained. On the test day, mice were habituated to the test cage and then removed. A cylindrical food pellet (∼2 cm) was buried 6 cm beneath fresh bedding in one corner, and the mouse was placed in the opposite corner. Latency to touch the food was recorded; if the food was not found within 10 min (600 s), the test was terminated and latency was recorded as 600 s.

#### Marble burying test

The test was performed in a clean box (40 cm × 40 cm) filled with 7 cm of bedding, with 16 glass marbles placed on the surface. Mice explored freely for 30 min. Marbles covered by at least 50% were counted as buried, and the number of moved and unburied marbles was recorded at 20 and 30 min.

#### Barnes maze

The Barnes maze consisted of a circular platform (1 m diameter) with 20 evenly spaced holes, with a dark escape box beneath one hole and a bright light source above the platform. The test included one adaptation day, four training days, and tests on days 5 and 9. During training, mice were placed in the center and released from a black box for 3 min, three times daily; the time to find the escape box was recorded, followed by 1 min of rest in the box, and the platform was cleaned and rotated after each trial. On test days 5 and 9, the escape box was removed, and mice explored freely for 90 s.

#### Rotarod

The rotarod device was equipped with a drop sensor triggered when mice fell from the rotating rod. Mice were tested on two consecutive days. On day 1, the rod accelerated linearly from 4 to 40 rpm over 5 min; on day 2, the rod accelerated linearly from 8 to 80 rpm over 5 min. Each day included two tests, with a 30-min interval between tests.

### Electrocorticography (ECoG) / electromyography (EMG) electrode implantation and signal acquisition

Mice were anesthetized and fixed in a stereotaxic frame. Recording screws were placed epidurally above the bilateral hippocampus (AP = -1.45 mm, ML = ±1.9 mm, relative to bregma), with reference/ground wires above the cerebellum. Two flexible EMG wires were inserted into the left and right neck muscles. Dental cement was used for fixation, and mice recovered for 7 days.

After 24 h of acclimation to the recording cable, 24-h Electrocorticography (ECoG) / electromyography (EMG) signals were continuously acquired using a tethered data acquisition system (Medusa, Nanjing, China) at 1000 Hz, with online band-pass filtering at 0.5–100 Hz, 50-Hz notch filtering, and synchronized video recording. Raw EDF signals were downsampled to 250 Hz and segmented into 60-min blocks. The first two ECoG channels from the first night segment were selected for core analysis. Signals were then subjected to 50-Hz harmonic multi-notch filtering, fourth-order Butterworth band-pass filtering (0.5 – 100 Hz), and Z-score normalization.

### ECoG preprocessing and spectral analysis

PSD was calculated using Welch’s method (256-point window, 128-point overlap, 256-point FFT). Absolute and relative power were calculated for δ (1–4 Hz), θ (4–8 Hz), α (8–13 Hz), β (13–30 Hz), and γ (30–80 Hz) bands. Relative power was normalized to total power (1–100 Hz). Magnitude-squared coherence was used to quantify linear dependence and frequency-specific synchronization between two ECoG channels. Coherence values were calculated as described previously using Welch’s periodogram method (256-point Hamming window, 50% overlap). ^13^ Mean coherence values within predefined frequency bands were calculated for between-group comparison.

Phase locking value (PLV) was used to quantify phase synchronization between the two cortical ECoG channels. Signals were band-pass filtered (fourth-order Butterworth) and instantaneous phase was extracted via the Hilbert transform. PLV was calculated for delta (1–4 Hz), theta (4–8 Hz), alpha (8–13 Hz), beta (13–30 Hz), and gamma (30–80 Hz) bands, with values from 0 to 1; higher values indicate stronger synchronization. Analysis was restricted to the first 120 s of each segment.

Phase–amplitude coupling (PAC) between theta phase and gamma amplitude was quantified using the modulation index (MI). For each channel, signals were band-pass filtered into theta (4– 8 Hz) and gamma (30–80 Hz) bands, and instantaneous theta phase and gamma amplitude were obtained via the Hilbert transform. Theta phase was divided into 18 bins over [−π, π], and mean gamma amplitude per bin was normalized to sum to 1; MI was then calculated from this distribution. A larger MI indicates stronger theta–gamma coupling. PAC was computed per channel, and the average MI across the two channels was used for group-level analysis.

### Sleep-stage classification

Data were divided into 2.5-s segments and classified into wakefulness, NREM sleep, and REM sleep using the open-source software Accusleep. Wakefulness was defined by low-amplitude, mixed-frequency EEG with variable or sustained EMG activity. NREM sleep was characterized by enhanced slow-wave activity and reduced EMG tone. REM sleep was characterized by theta-dominant EEG and very low-neck muscle EMG activity. Automatic classification used a trained model with manual annotation of a subset of segments as reference. The percentage of valid recording time occupied by each state was calculated separately for the 12-h light and dark periods. Between-group comparisons used mouse-level values.

### EEG analysis during social interaction

EEG, video, and social interaction were synchronized. Mice underwent 1-min social contact sessions separated by 10 min, with a new same-sex stranger mouse each time. Nonsocial behavior was defined as mice being far apart. Social contact start was defined as the moment mice began to approach and were about to touch. Active social interaction was defined as the experimental mouse sniffing the anogenital region of the stimulus mouse. A sequence from nonsocial behavior to social contact start to active social interaction, without interruption and with social interaction >3 s, was defined as an effective social behavior. The first three effective social behaviors per trial were selected. Social contact start was set as the anchor (0 s). Continuous EEG from −2.5 to +2.5 s around the anchor was extracted and band-pass filtered at 0.5–100 Hz with 50-Hz notch filtering. PSD was calculated separately for Pre (−2.5–0 s) and Post (0–2.5 s) using Welch’s method. Total power (0.5–100 Hz), absolute power, and relative power were calculated for δ (1–4 Hz), θ (4–8 Hz), α (8–13 Hz), β (13–30 Hz), and γ (30–80 Hz). Pre/Post ratio and normalized change (Post − Pre)/(Pre + Post) were also calculated. For theta and gamma, signals from −2.5 to +2.5 s were band-pass filtered with a fourth-order Butterworth filter, and power time series were calculated using a 0.2-s window with a 0.02-s step. A single social contact event was used as the observational unit.

### Western blotting

Protein samples were lysed in RIPA buffer containing protease inhibitor cocktail, centrifuged (10,000 × g, 10 min, 4 °C), and supernatants were heated at 100 °C for 30 min. Samples were separated by SDS–PAGE (Tris-Gly 4–20%, 15-well) at 80 V for stacking and 120 V for separating gels, and transferred to PVDF membranes at 200 mA. Membranes were blocked with 5% BSA in TBS-T for 1 h and incubated overnight at 4 °C with primary antibodies: Tbr1 (1:1000, Abcam, ab183032) and GAPDH (1:3000, GB11002, Service bio). After TBS-T washes, membranes were incubated with secondary antibodies, and bands were visualized using chemiluminescent substrate (Thermo Fisher, 32106).

### Immunofluorescence

Five-month-old mice were deeply anesthetized with isoflurane and transcranial perfused with cold PBS and 4% PFA. Brains were post-fixed overnight at 4 °C, embedded in agarose, and coronally sectioned at 50 μm. Sections were blocked with 5% BSA and 0.1% Triton X-100 for 120 min and incubated overnight at 4 °C with primary antibodies: Ctip2 (1:500, Abcam, ab18465), Tbr1 (1:500, Abcam, ab183032), PV (1:1000, Abcam, ab181086), and IBA1 (1:1000, Wako, Cat# 019-19741, RRID: AB_839504). After PBS washes, sections were incubated with fluorescent secondary antibodies and DAPI for 1 h at room temperature. Sections from different groups were distributed across staining batches with identical reagents and imaging settings. Quantification was performed in anatomically matched ROIs in ImageJ. TBR1+ cells were counted in RSG, RSA, PtA, and S1Tr (700 × 250 μm for RSG/RSA; 700 × 300 μm for PtA/S1Tr) and in the amygdala (300 × 300 μm). Cortical neurons were counted in S1Tr layer 5 (650–850 μm; 500 × 200 μm) for Ctip2+ and TBR1+, and layer 6 (850–1350 μm; 200 × 300 μm) for TBR1+. PV+ cells were counted in RSA and RSG (700 × 250 μm), and IBA1+ cells in RSG, RSA, and PtA (200 × 200 μm).

### Statistical analysis

Data are presented as mean ± SEM, with n indicated in figure legends. Two-group comparisons used two-tailed unpaired t tests with Welch’s correction. For three or more groups, one-way or ordinary two-way ANOVA was used according to the experimental design, followed by Tukey’s, Dunnett’s, or Šídák’s correction for multiple comparisons. Correlation analysis was performed using two-tailed Pearson correlation. p < 0.05 was considered statistically significant. Analyses were performed using GraphPad Prism.

## Data and code availability

The raw data required to reproduce these findings will be made available by the corresponding author upon request.

## Acknowledgments

This work was supported by the following grants: National Science and Technology Major Project (2025ZD0214700), National Natural Science Foundation of China (82430046 to Z.Q.), and Project of Medical Technology Research and Transformation supported by Shanghai Municipal Health Commission (2024ZZ1007).

ChatGPT (OpenAI) was used to assist with language editing, manuscript organization, and drafting. The authors independently reviewed and verified all scientific content, analyses, interpretations, and references and accept full responsibility for the manuscript.

## Author contributions

Z.Q., T.L., and Z.J. conceived and designed the study. T.L. performed experiments with assistance from X.W. and C.Y., Y.Y., and Y.Z.. Z.Q. supervised the project and wrote the manuscript.

## Supplemental information

Supplemental information can be found online.

