## Supplemental figures for "AAV-mediated TBR1 replacement ameliorates behavioral and electrophysiological abnormalities in *Tbr1*-haploinsufficient mice"

### Supplemental Figures 1-4

Supplementary Figure 1

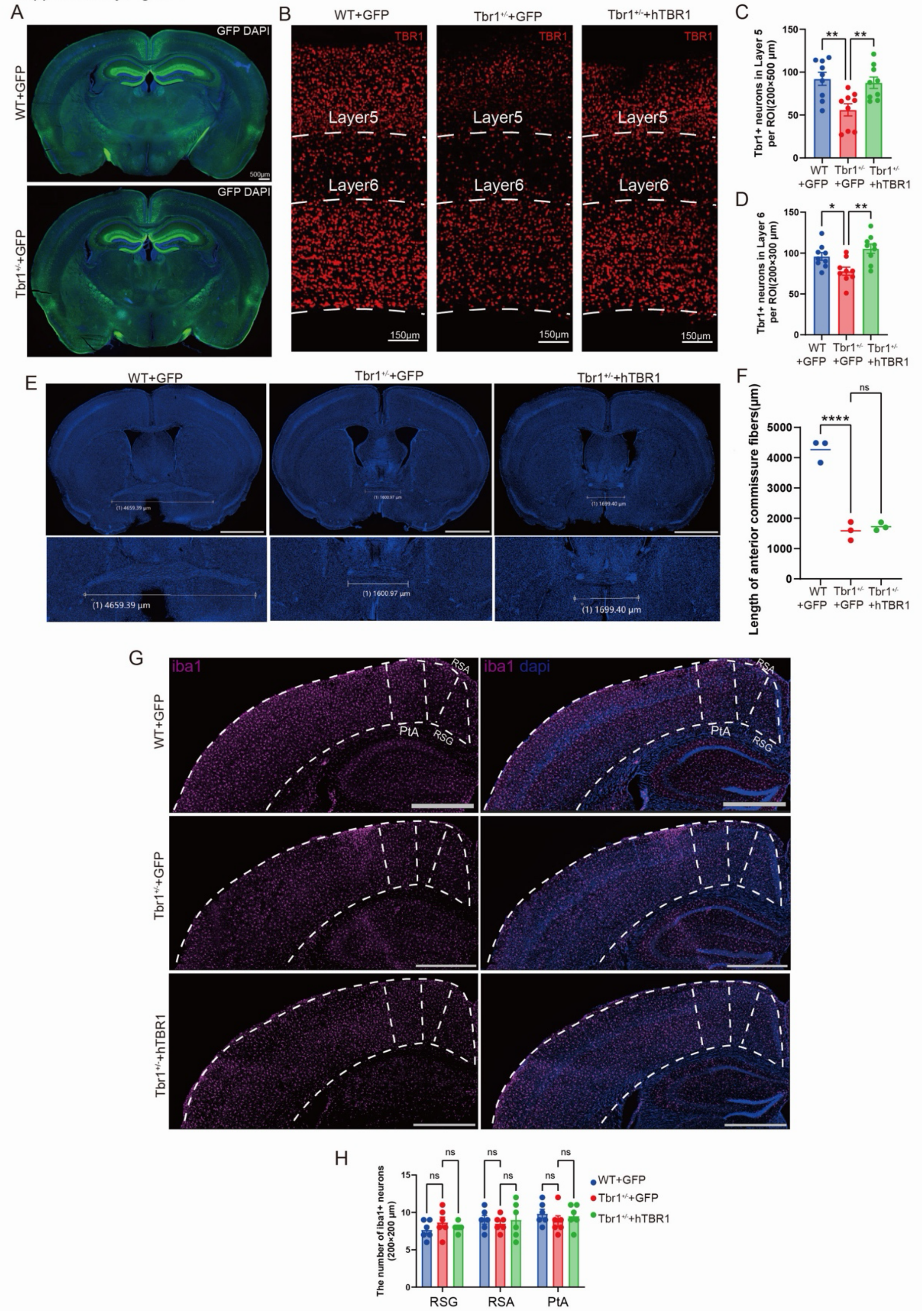

**Figure S1. Cortical TBR1 expression, anterior commissure morphology, and IBA1-positive cell counts following neonatal AAV-mediated TBR1 replacement**

Analyses were performed in wild-type mice receiving control AAV-EGFP (WT+GFP), *Tbr1*$^{+/-}$ mice receiving AAV-EGFP (*Tbr1*$^{+/-}$+GFP), and *Tbr1*$^{+/-}$ mice receiving AAV-hTBR1 (*Tbr1*$^{+/-}$+hTBR1).

(A) Immunofluorescence images showing the expression of GFP in 5-month-old WT and Tbr1+/- mice following AAV-PHP.eB-hSyn-EGFP injection at Postnatal Day 3, with DAPI counterstaining. Scale bars, 500μm.

(B) Representative immunofluorescence images showing TBR1 expression (red) in cortical layers V and VI of the S1Tr. boundaries. Dashed lines indicate cortical layer boundaries.

(C and D) Quantification of TBR1-positive cells in layer VI per 200 × 300 μm region of interest (ROI;D) and layer V per 200 × 500 μm ROI (C). Both layer V and layer VI were analyzed in the S1Tr. Each point represents one section (three sections per mouse; n = 3 mice per group). Data were analyzed using ordinary one-way ANOVA followed by Dunnett's multiple comparisons test.

(E) Representative coronal brain sections showing the anterior commissure (upper), with enlarged views below. Horizontal measurement lines indicate the quantified mediolateral extent of the anterior commissure. Sections were counterstained with DAPI. Scale bars, 2mm.

(F) Quantification of the anterior commissure extent measured as illustrated in E. Each point represents an individual mouse, with n = 3 mice per group. Data were analyzed using ordinary one-way ANOVA followed by Dunnett's multiple comparisons test.

(G) Representative immunofluorescence images showing IBA1-positive cells (magenta; left) and merged IBA1 and DAPI staining (blue; right). Dashed outlines delineate the indicated cortical regions. RSG, granular retrosplenial cortex; RSA, agranular retrosplenial cortex; PtA, parietal association cortex. Scale bars, 1mm.

(H) Quantification of IBA1-positive cells per 200 × 200 μm ROI in the RSG, RSA, and PtA. Each point represents one section (three sections per mouse; n = 3 mice per group). Data were analyzed using ordinary two-way ANOVA followed by Dunnett's multiple comparisons test.

Scale bars, 500 μm (A); 150 μm (B); 2mm(E); 1mm(G) Bars represent mean ±SEM; horizontal lines in F indicate mean. Both male and female mice were included Asterisks indicate significance for the comparisons marked by brackets: *$p < 0.05$, **$p < 0.01$, and ****$p < 0.0001$; ns, not significant.

Supplementary Figure 2

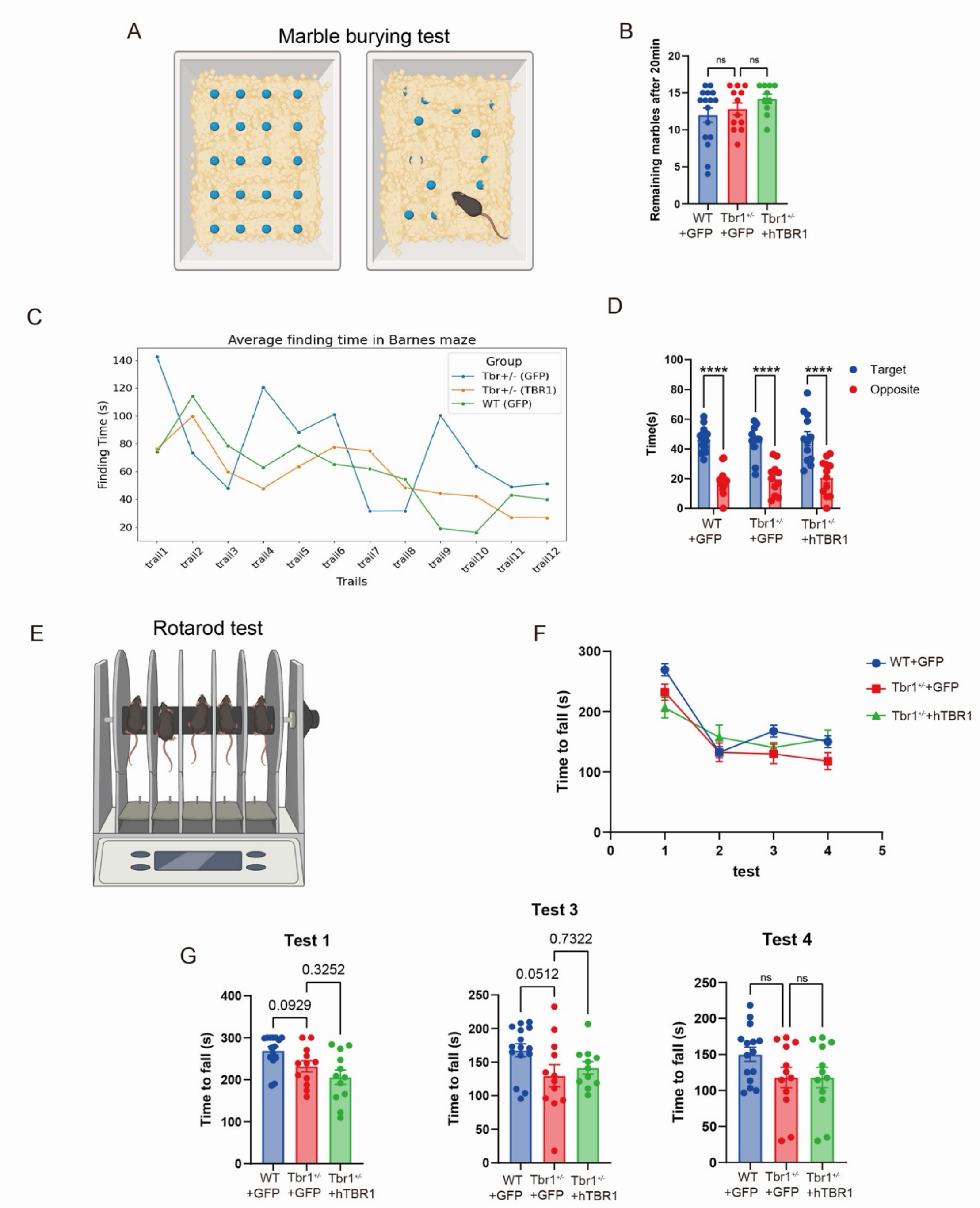


**Figure S2. Marble burying, Barnes maze, and rotarod performance following neonatal AAV-mediated TBR1 replacement**

Behavioral assessments were performed in wild-type mice receiving control AAV-EGFP (WT+GFP), *Tbr1*$^{+/-}$ mice receiving AAV-EGFP (*Tbr1*$^{+/-}$+GFP), and *Tbr1*$^{+/-}$ mice receiving AAV-hTBR1 (*Tbr1*$^{+/-}$+hTBR1).

(A) Schematic of the marble-burying test, illustrating the initial arrangement of marbles and their distribution following testing.

(B) Number of marbles remaining unburied after the 20-min test. WT+GFP, n=16; Tbr1$^{+/-}$+GFP, n=12; Tbr1$^{+/-}$+hTBR1, n=10. Data were analyzed using ordinary one-way ANOVA followed by Dunnett's multiple comparisons test.

(C) Mean target-finding latency across 12 training trials in the Barnes maze.

(D) Time spent in the target and opposite regions of the Barnes maze. Blue and red indicate the target and opposite regions, respectively. Brackets indicate comparisons between regions within each experimental group. The test phase comprised a 1-day habituation period, 4 training days (three trials per day), and probe tests on days 5 and 9. WT+GFP, n=16; Tbr1$^{+/-}$+GFP, n=11; Tbr1$^{+/-}$+hTBR1, n=12. Data were analyzed using ordinary two-way ANOVA followed by Šídák's multiple comparisons test.

(E) Schematic of the accelerating rotarod protocol. Rotation speed increased from 4 to 80 rpm over 12 min. Mice underwent four trials separated by 30-min intervals.

(F) Latency to fall across the four rotarod trials. WT+GFP, n=15; Tbr1$^{+/-}$+GFP, n=12; Tbr1$^{+/-}$+hTBR1, n=12.

(G) Individual values and group summaries for latency to fall during rotarod trials 1, 3, and 4. WT+GFP, n=15; Tbr1$^{+/-}$+GFP, n=12; Tbr1$^{+/-}$+hTBR1, n=12. Data were analyzed using ordinary one-way ANOVA followed by Dunnett's multiple comparisons test.

Panel C shows group means. Data in B, D, F, and G are presented as mean ±SEM. Each point in B, D, and G represent an individual mouse. Both male and female mice were included. ****$p < 0.0001$; ns, not significant. Exact p values are shown where indicated.

Supplementary Figure 3

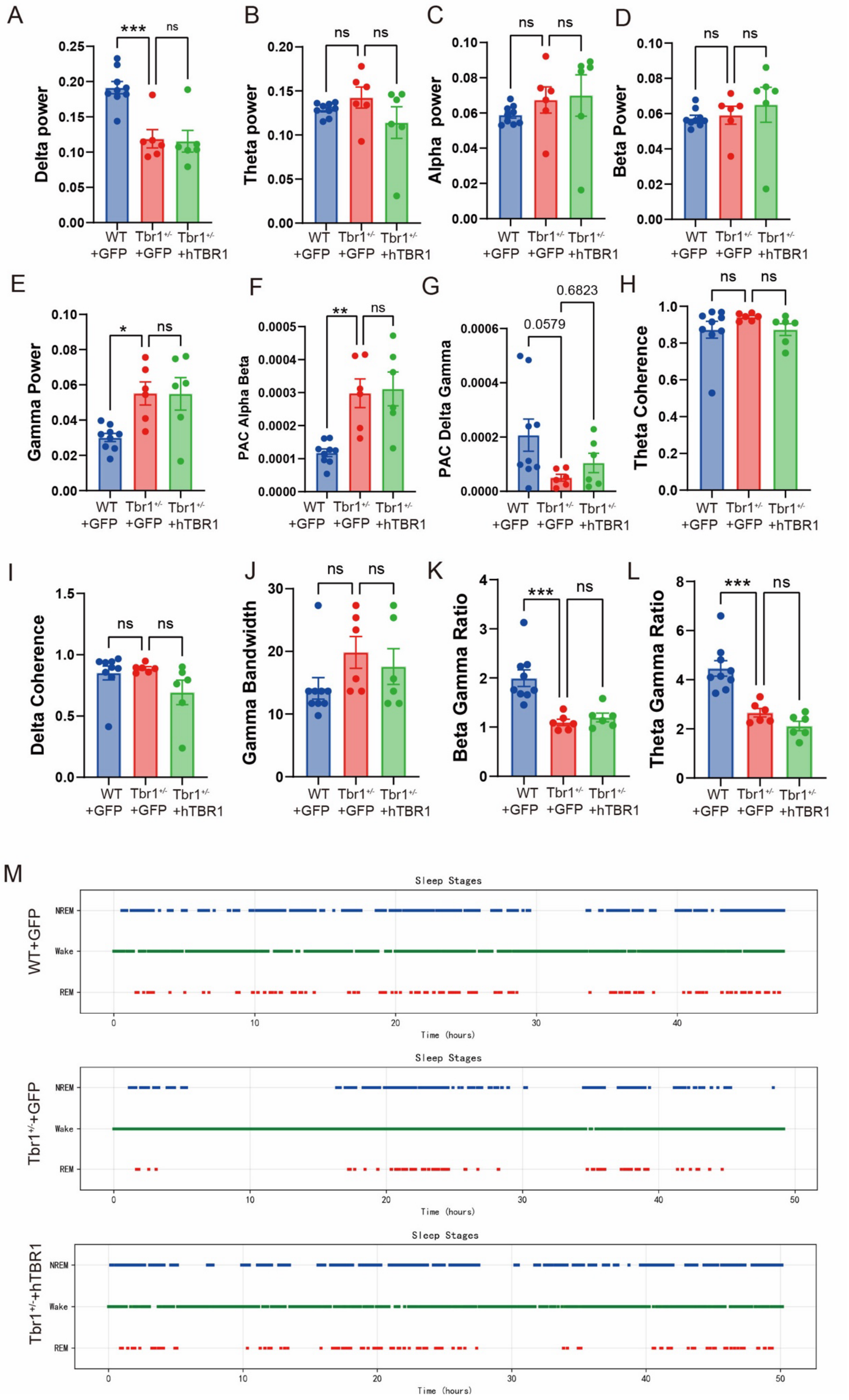

**Figure S3. Additional EEG spectral, coupling, and sleep–wake analyses following neonatal AAV-mediated TBR1 replacement**

Analyses were performed in wild-type mice receiving control AAV-EGFP (WT+GFP), *Tbr1*$^{+/-}$ mice receiving AAV-EGFP (*Tbr1*$^{+/-}$+GFP), and *Tbr1*$^{+/-}$ mice receiving AAV-hTBR1 (*Tbr1*$^{+/-}$+hTBR1).

(A–E) Quantification of electroencephalographic (EEG) power in the delta (A), theta (B), alpha (C), beta (D), and gamma (E) frequency bands. Power spectral density was computed using Welch's method (256-point window, 128-point overlap, 256-point FFT), and absolute power ($\mu V^2$) was analyzed for each frequency band. (F and G) Alpha–beta (F) and delta–gamma (G) phase–amplitude coupling (PAC). PAC was quantified using the modulation index (MI), with alpha (8–13 Hz) and delta (1–4 Hz) as the phase-providing bands and beta (13–30 Hz) and gamma (30–80 Hz) as the amplitude-providing bands, respectively. (H and I) EEG coherence in the theta (H) and delta (I) frequency bands between all analyzed channel pairs.

(J) Gamma bandwidth. Gamma bandwidth was defined as the frequency range of the gamma peak in the power spectrum, calculated from the power spectral density and expressed in Hz. (K and L) Beta-to-gamma (K) and theta-to-gamma (L) power ratios.

(M) Temporal distribution of wakefulness, non-rapid eye movement (NREM) sleep, and rapid eye movement (REM) sleep in the three experimental groups. Green, blue, and red marks indicate wakefulness, NREM sleep, and REM sleep, respectively. Each timeline represents an individual mouse, with a recording duration of 48 h and a 2.5-s scoring epoch length.

EEG measures in A–L were calculated from the 24-h continuous recording period across all vigilance states, using the following frequency-band boundaries: delta (1–4 Hz), theta (4–8 Hz), alpha (8–13 Hz), beta (13–30 Hz), and gamma (30–80 Hz).. Bar graphs show mean ±SEM. Each point represents an individual mouse. WT+GFP, n=9; Tbr1$^{+/-}$+GFP, n=6; Tbr1$^{+/-}$+hTBR1, n=6. Statistical analyses were performed using ordinary one-way ANOVA followed by Dunnett's multiple comparisons test. Both male and female mice were included. Asterisks indicate significance for the comparisons marked by brackets: *p < 0.05, **p < 0.01, and ***p < 0.001; ns, not significant.

Supplementary Figure 4

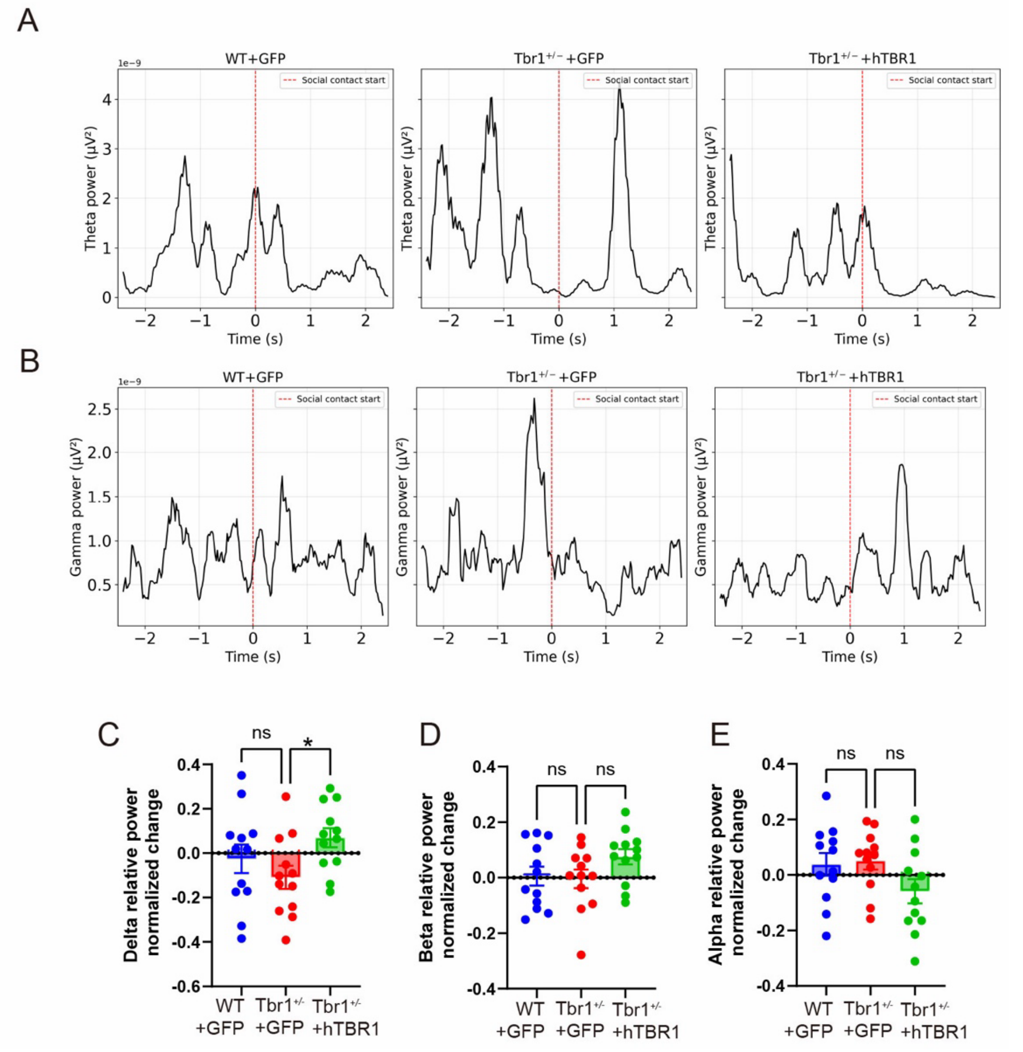


**Figure S4. EEG power dynamics during social interaction following neonatal AAV-mediated TBR1 replacement**

Analyses were performed in wild-type mice receiving control AAV-EGFP (WT+GFP), *Tbr1*$^{+/-}$ mice receiving AAV-EGFP (*Tbr1*$^{+/-}$+GFP), and *Tbr1*$^{+/-}$ mice receiving AAV-hTBR1 (*Tbr1*$^{+/-}$+hTBR1).

(A and B) Theta (A) and gamma (B) power time courses aligned to social-contact onset in the three experimental groups. Traces span 2.5 s before and 2.5 s after contact. Red dashed lines indicate social-contact onset (time 0). Each trace represents a single social-interaction event. (C–E) Normalized changes in relative delta (C), beta (D), and alpha (E) power associated with the transition from nonsocial behavior to active social interaction. Horizontal lines at zero indicate no change. Relative power was calculated as the power in each frequency band divided by the total power in the 1–100 Hz range (expressed as a percentage). Normalized changes in C–E were calculated as (social − nonsocial) / (social + nonsocial). Each point represents a social-interaction event, with 12 events analyzed per group (n = 3 WT+GFP, 2 $Tbr1^{+/-}$+GFP, and 3 $Tbr1^{+/-}$+hTBR1 mice). Data were analyzed using ordinary one-way ANOVA followed by Dunnett's multiple comparisons test.

Frequency bands were defined as delta (1–4 Hz), theta (4–8 Hz), alpha (8–13 Hz), beta (13–30 Hz), and gamma (30–80 Hz). Bar graphs show mean ±SEM. All mice were female. *$p < 0.05$ for the comparison indicated by the bracket; ns, not significant.
